## Supplementary Information for "scMD: cell type deconvolution using single-cell DNA methylation references"

### Supplementary Figures

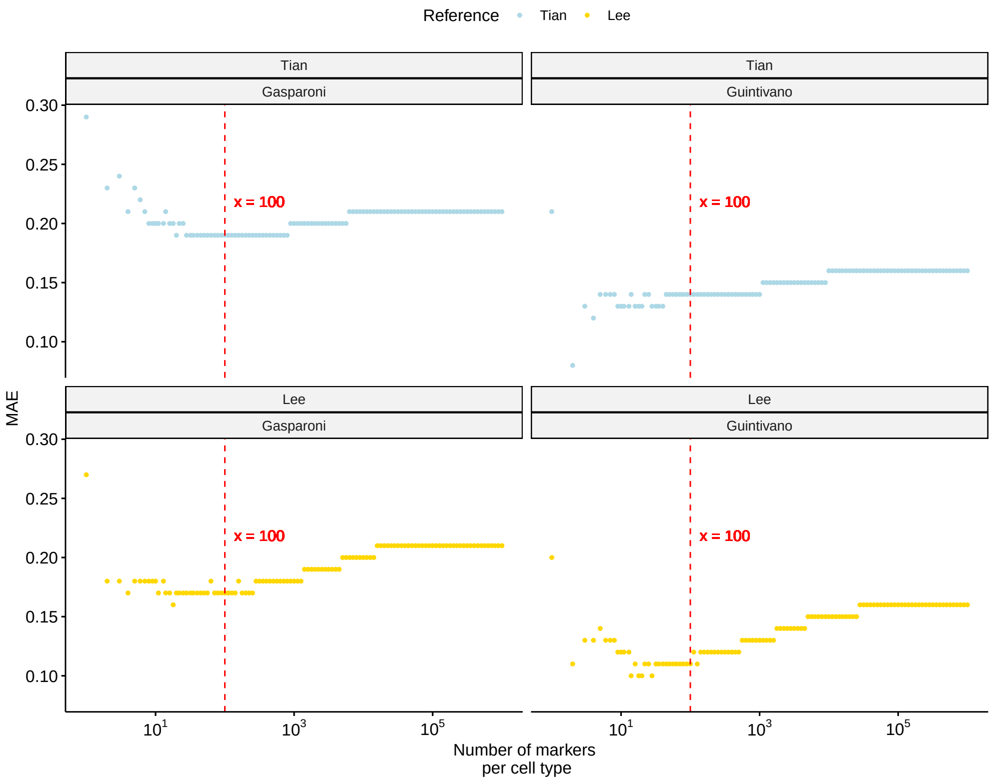


Figure S1: The mean absolute error (MAE) between estimated DNAm fraction and measured DNAm fraction as a function of different numbers of markers per cell type. Each row panel is a reference dataset, and each column panel is a bulk dataset.

### Supplementary Tables

Table S1: Number of DNAm sites of scDNAm datasets during the process before generating the signatures. The cluster-level CpGs are matched to bulk data and resulted in 450k, 850k, or 25 million (for WGBS) CpGs.

| | single-cell CG+CH | single-cell CG | cluster-level CG |
|---|---|---|---|
| Lee et al., 2019 (mean (sd)) | 48,216,050 (49,294,600) | 2,312,785 (2,374,714) | 53,009,685 (53,407,452) |
| Tian et al., 2022 (mean (sd)) | NA | NA | 53,601,064 (53,739,700) |

Table S2: Summary of sorted-cell and bulk datasets for validation. The first three datasets are based on sorted cells, while the rest are bulk datasets.

| Dataset | Technical platform | Cell types (Sample size) |
|---|---|---|
| Mendizabal et al., 2019 | WGBS | neurons (25) and oligodendrocyte (20) |
| Guintivano et al., 2013 | Illumina 450k | neurons (29) and non-neurons (29) |
| Gasparoni et al., 2018 | Illumina 450k | neurons (31) and non-neurons (31) |
| ROS (Religious Orders Study) | Illumina 450k | Bulk (49) De Jager et al., 2014 |
| NAc (Nucleus Accumbens) | Illumina 850k & RNA-seq | Bulk (211) Markunas et al., 2021 |
| MSBB (Mount Sinai Brain Bank) | Illumina 850k | Bulk (201) M. Wang et al., 2018 |

Table S3: Cell-type-specific (CTS) differential fraction analyses of different phenotypes in MSBB bulk data using scMD and EpiSCORE estimated fractions. Correlation coefficients and p-values for phenotype-fraction associations in different cell types are provided.

| | | Correlation (p-value) | | | | | | | |
|---|---|---|---|---|---|---|---|---|---|
| Phenotype | Method | Astro | Micro | Endo | Oligo | OPC | Inh | Exc | Neuron |
| Age | scMD | 0.097 (0.172) | 0.127 (0.072) | -0.012 (0.866) | 0.058 (0.413) | -0.173 (0.014) | -0.145 (0.040) | -0.057 (0.424) | -0.091 (0.198) |
| | EpiSCORE | -0.030 (0.675) | 0.048 (0.501) | 0.065 (0.357) | 0.104 (0.140) | -0.059 (0.409) | | | -0.047 (0.506) |
| CDR | scMD | 0.013 (0.853) | 0.217 (0.002) | -0.073 (0.306) | 0.215 (0.002) | -0.198 (0.005) | -0.142 (0.044) | -0.284 (0.000) | -0.255 (0.0003) |
| | EpiSCORE | 0.150 (0.034) | 0.138 (0.051) | 0.039 (0.579) | 0.236 (0.001) | -0.095 (0.180) | | | -0.207 (0.003) |
| CERAD | scMD | 0.019 (0.788) | 0.035 (0.624) | 0.010 (0.893) | -0.041 (0.561) | 0.037 (0.598) | 0.013 (0.855) | 0.0180 (0.804) | 0.017 (0.808) |
| | EpiSCORE | 0.106 (0.244) | -0.018 (0.799) | 0.015 (0.833) | 0.052 (0.466) | -0.114 (0.107) | | | 0.070 (0.325) |
| Braak | scMD | 0.057 (0.530) | 0.189 (0.036) | -0.079 (0.387) | 0.142 (0.118) | -0.037 (0.688) | -0.106 (0.243) | -0.258 (0.004) | -0.195 (0.005) |
| | EpiSCORE | 0.063 (0.374) | 0.070 (0.327) | 0.134 (0.059) | 0.233 (0.001) | -0.145 (0.040) | | | -0.172 (0.015) |

Table S4: Number of identified differentially methylated cytosines (DMCs) in individual cell types using CellDMC and scMD estimated cellular fractions, with FDR < 0.05. We adjusted for covariates of sex and race.

| | Number of DMCs | | | | | | |
|---|---|---|---|---|---|---|---|
| | Astro | Micro | Endo | Oligo | OPC | Inh | Exc |
| Age | 7 | 38 | 1 | | 13 | 8 | |
| CDR | 221 | | 2 | | 20 | | |
| CERAD | | | | 13 | | 69 | 23 |
| Braak | | | 5 | | | 17 | |
